# The Enemy of my Enemy is my Friend: Immune-Mediated Facilitation Contributes to Fitness of Co-Infecting Helminths

**DOI:** 10.1101/2021.08.25.457665

**Authors:** Francesca Dagostin, Chiara M. Vanalli, Brian Boag, Renato Casagrandi, Marino Gatto, Lorenzo Mari, Isabella M. Cattadori

## Abstract

Our conceptual understanding of immune-mediated interactions between parasites is rooted in the theory of community ecology. One of the limitations of this approach is that most of the theory and empirical evidence has focused on resource or immune-mediated parasite competition and yet, there is ample evidence of positive interactions between species that could be generated by immune-mediated facilitation. Here, we develop an immuno-epidemiological framework and apply it to longitudinal infection data of two gastrointestinal helminths that infect a population of free-living rabbits to investigate, through model testing, the mechanisms of immune-mediated facilitation in dual infections. Simulations show that weakened, species-specific IgA antibody responses and unequal, albeit low, IgA cross-reactions explain higher parasite intensities in dual compared to single infections, for both helminths. Simulations also show that rabbits with dual infections shed more free-living stages that survive fort longer in the environment, implying greater onward transmission than hosts with single infections. These findings support the hypothesis that the two helminths interact through immune-mediated facilitation which contributes to greater fitness and the long-term co-circulation of both species in the host population.

## Introduction

Interference between parasite species, mediated via the host’s immune response, is one of the processes frequently proposed to explain interactions between different parasite species, or genetically diverse strains, that co-infect the same host [1–6]. In these instances, immunity is expected to preferentially target the more abundant or virulent parasite and, by reducing its intensity, attenuate the competition on the less abundant species. The two species can still co-exist, but their relative fitness depends on the net outcome of these interactions.

An alternative scenario to competition is where immune-mediated interactions lead to facilitation of either one or both of the co-infecting species [7] (figure 1). For example, many parasites can suppress or divert the immune response in favour of their own survival [8], and this action can benefit a second parasite species through a bystander effect [9,10]. Similarly, because of the polarization and function of different branches of the immune system, the response developed against one species can reduce or prevent the reaction against a co-infecting species [11–14]. In these interactions, the net benefit obtained from co-infection is asymmetrical (figure 1*c*). The facilitator parasite can control the immune response, or become its target, and in so doing facilitates the second parasite species that profits more than the facilitator, and consequently has greater fitness than when it is the sole parasite in the host. Symmetrical immune-mediated facilitation emerges when both parasite species benefit from their co-infections (figure 1*b*). This can occur through the reduction, suppression or evasion of the immune response against each species, which gains both through greater vital rates and abundance [15–17]. Immune tolerance could be considered an extreme example of facilitation since the host immunity is engaged in repairing the injuries caused by the parasites rather than controlling the infections [18]. Importantly, under these scenarios a parasite will have a selective advantage when it infects hosts already parasitized with the other species, so long as the facilitation is not so strong as to destabilize the interactions [19] or reduce host survival [20,21].

**Figure 1.**
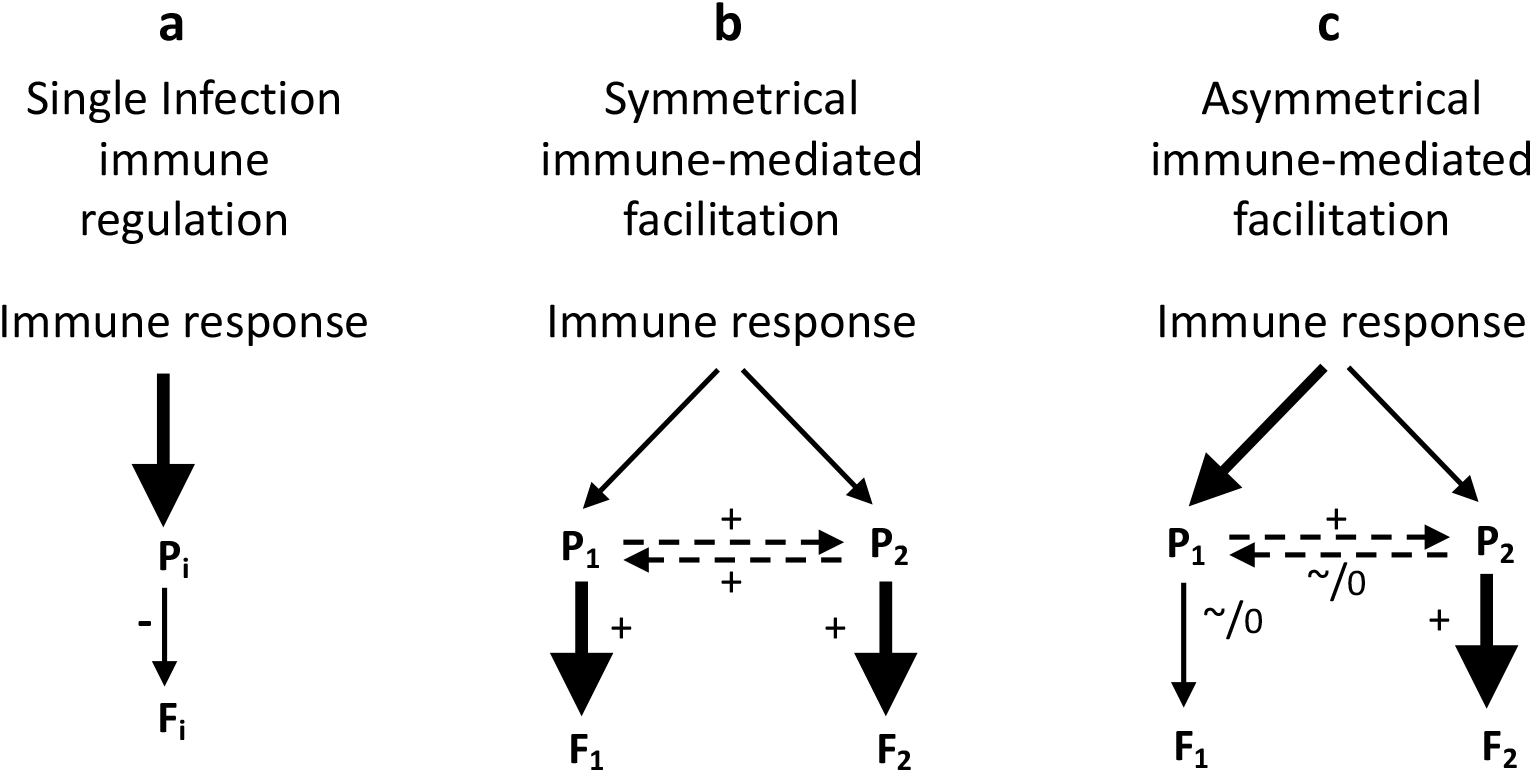
Scenarios of immune-parasite interaction in single infections (a) and dual-infections (b and c) for two parasite species (P_i_, i= 1, 2) and consequences for their fitness (F_i_, i= 1, 2). Symmetrical and asymmetrical immune-mediated facilitation is presented for the dual infection. We report: the magnitude (arrow thickness) and type of the effect (null= 0, sub-typical= ~, positive= +, negative= -) and the parasite indirect interactions (dotted arrow). **a-** Standard scenario were immunity directly affects parasite, P_i_, traits (i.e. abundance, development or fecundity) and reduces its fitness, f_i_; examples: many helminth species, including *T. retortaeformis* in rabbits (Cattadori et al. 2005). **b-** Immunity decreases against each parasite, P_1_ and P_2_, and this benefits their fitness, F_1_ and F_2_, compared to case *a*; P_1_-P_2_ interactions (e.g. positive or unclear reaction) do not reduce the positive net immune effect on F_1_ and F_2_; examples: hookworm and *Ascaris lumbricoides* in humans (Fleming et al. 2006), HIV-*Mycobacterium tuberculosis* in humans (Pawlowski et al. 2012), this study. **c-** Processes and impacts are as described in case *b*, however, P_1_-P_2_ asymmetrical interactions benefit the fitness F_2_ of P_2_ but lead to sub-typical (~) or unclear (0) increase in fitness F_1_ for P_1_, compared to case *a*; example: *Trichuris muris-Schistosoma mansoni* in mice (Bickle et al. 2008) or *H. polygyrus-Trichinella spiralis* in mice (Behnke et al. 1993).

From an ecological perspective, the extent of immune-mediated parasite interaction is the outcome of two fundamental components. First, the immune response against one parasite species that is diverted to fight other parasites, e.g. the loss of specific antibodies that cross-react with other infections, and second, the immune constraints this parasite species experiences by the presence of other species, e.g. the effect of cross-reacting antibodies from other infections (figure 1*b,c*). Therefore, a positive net impact for each species, including any parasite manipulation of the immune reactions, would be expected to facilitate co-infections both within the host and among the co-circulating parasites at the host population level. Theory has shown that in the absence of strong competition between two parasite species, a weak immune-mediated symmetrical facilitation increases the virulence/abundance of both parasites, however, under stronger facilitation parasite growth escapes regulation and the system becomes destabilized [19].

To test for immune-mediated interactions leading to facilitation, we applied an immuno-epidemiological model to 23 years of infection data and asked whether the interaction between two helminths in dual infected hosts could be explained by immune facilitation, and whether there were positive consequences for parasite fitness, or onwards transmission, at the host population level. We used two helminth species, *Trichostrongylus retortaeformis* and *Graphidium strigosum*, that inhabit different gastrointestinal niches of the European rabbit (*Oryctolagus cuniculus*), the small intestine and stomach respectively, implying that they were unlikely to exhibit direct competition. While both helminths cause chronic infections, *T. retortaeformis*, and less so *G. strigosum*, shows evidence of regulation of abundance and fecundity through host immunity [22–27]. Interestingly, field studies found higher intensities of the first and to a lesser extent the second in rabbits with both helminths, when compared to hosts with single infections [28,29], suggesting a possible positive interaction between the two species.

We investigated two hypotheses within the processes of immune regulation. If symmetrical facilitation is important, we should expect both helminth species from dual infections to experience reduced immune constraints and higher abundance and/or production of free-living stages, than parasites from single infections (figure 1*b*). However, if asymmetrical facilitation drives the interaction between the two species, then the dynamics for the facilitated parasite should be enhanced but the facilitator should not show substantial changes, when compared to single infections (figure 1*c*). In the opposite scenario of reduced dynamics, such as decreased intensity of infection and/or parasite fitness for one of both species, this should support immune-mediated interference.

We tested different candidate frameworks, and model selection was based on the ability of each model to describe the observed dynamics of infection, while identifying a parsimonious mechanism of host-parasite and parasite-parasite interactions. Since we are interested in processes of facilitation through host immunity, rather than competition, and given that the two helminths inhabit different gastrointestinal niches, parasite interference for host resources, including manipulation of host’s metabolism or the interaction with the gut microbiome, were beyond the scope of this study.

## Material and Methods

### The System

Rabbits become chronically infected with the two helminths by eating herbage contaminated with infective larvae; once in the host, larvae develop into adults that shed eggs with the hosts’ faeces (figure S1). Laboratory experiments showed that rabbits develop a type 2 anti-inflammatory response against both species. However, while this regulates both *T. retortaeformis* intensity and body growth, which then affects fecundity, it appears to have a weaker effect against *G. strigosum* that maintains high intensities throughout the infection [24–27]. A broader investigation of the immune profile in experiments of infection and anthelminthic treatment found a down-regulation of genes expressing type 1, type 2 and T-regulatory responses during reinfections with *G. strigosum* and much less so with *T. retortaeformis*, both in single and dual infections [23]. There was no evidence that *G. strigosum* contributed to this reduction, suggesting that the host rather than the parasite was probably responsible for the observed immune down-regulation [23]. The general conclusions that *T. retortaeformis*, and to a lesser extent *G. strigosum*, is affected by host immunity were also proposed in field studies [22,28-31].

The dynamics of the two helminths could also be negatively affected by ecological processes of intra-specific competition for resources. Previous studies indicated that this is probably not the main mechanism of regulation for *T. retortaeformis* but could play a role for *G. strigosum*, especially at high abundances [26,31]. For example, rabbits nutritionally constrained, through coprophagic restriction and on a fixed diet, carried the same *T. retortaeformis* intensities as animals on a fixed diet only [32].

An important component of soil-transmitted helminths is the development and survival of eggs and larvae in the environment. Temperature and rainfall affect the survival of both *T. retortaeformis* and *G. strigosum* free-living stages [33–35] while at the host population level, climate and seasonality were found to be important for subsequent infection and thus parasite fitness [31].

### Model Datasets

Here, we present the datasets and assumptions used for our mechanistic immuno-epidemiological model, while in the next section we describe the model framework. The study focused on a population of rabbits (population A) sampled monthly between 1980 and 2002 from our site in Scotland. For every rabbit collected, the abundance of *T. retortaeformis* and *G. strigosum* was quantified by aliquots using standardized parasitological techniques [36]. We classified rabbits with one helminth species as single-infected and with both species as dual-infected. It is possible that some of the adults with single infections may have been previously infected with the other species that was subsequently eliminated. Rabbits do not appear to be able to clear *G. strigosum* [23-25,28,37] but old hosts can remove *T. retortaeformis* [24,27], although in endemic settings reinfection is relatively fast and the fraction of single infected hosts is really quite low; nevertheless, dubious cases were removed. To avoid interference with myxoma virus infection [11,28] rabbits with myxomatosis symptoms were excluded. Host age was arranged into eight classes, from age class 1 (one-month old) to age class 8 (8+ months old) [22,28] and lifespan was considered to be about one year [38].

Population A provides 23 years of robust individual host data on the two helminths but lacks information on host immunity. Since our aim is to integrate the immune response of the rabbit with the dynamics of its parasites, we used data on serum antibody IgA and IgG data obtained from a second rabbit population (population B) located ~5 kilometres away in a similar ecosystem and sampled for a much shorter time period (2004-2011), following the same standardized procedures as in population A [29]. Antibody data were available for six out of those eight years. Given its shorter time series of infection and following a preliminary investigation (SI-1.3 and 1.4), antibodies from population B were used to inform the parameterization of the immune response for the longer sampling of population A. This was based on similar dynamics of the two helminths (compare figure S2 with figure 2 and statistical results in SI-1.3 and 3.3) and the rabbits’ age structure (SI-1.4) at the two populations.

**Figure 2.**
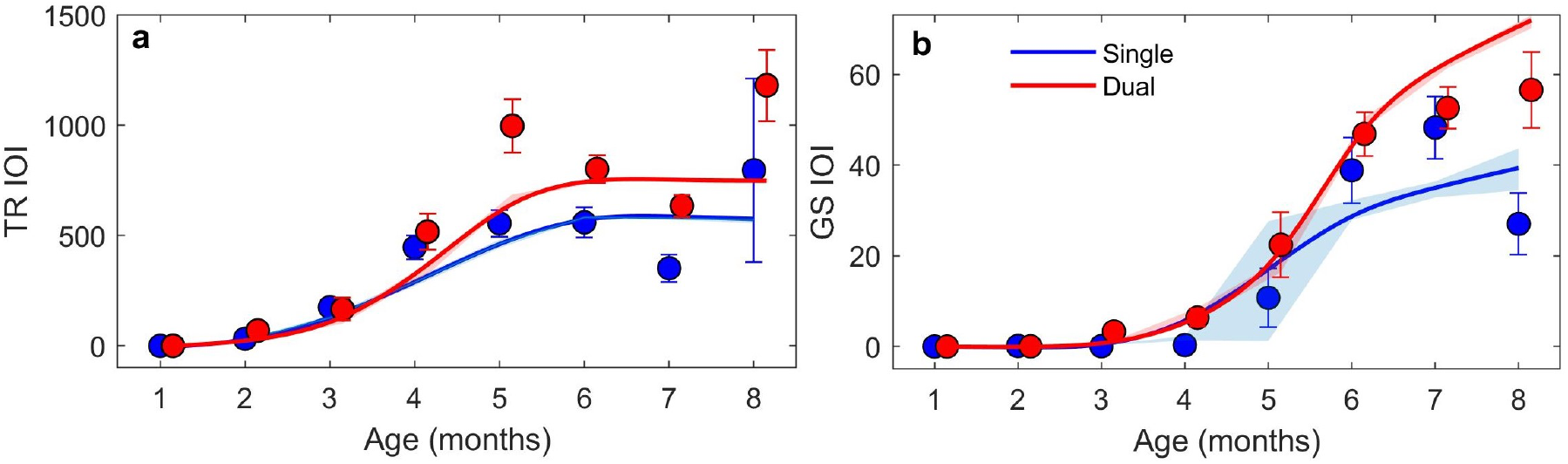
Relationship between intensity of infection (IOI) and host age for *T. retortaeformis* (TR, a) and *G. strigosum* (GS, b) in single- (blue) and dual- (red) infected hosts from population A. Mean and S.E. data are reported from individual-based model simulations (lines and shadow bands) and field monitoring (circles and bars, these latter ones calculated under the assumption that the data follow a negative binomial distribution). Only simulations referring to collected rabbit data are reported. Small S.E. bars and bands could be masked by circles or lines. IOI at age 1 is forced to start at 0 since rabbits are exposed to the risk of infection at about 30 days of age when they switch from milk to herbage.

We selected species-specific IgA antibodies to represent host immunity against each of the two helminths. Naturally, this is an over-simplification of the immune response, and it was determined by the following reasons. First, IgA is the most abundant immunoglobulin at mucosal surfaces, including the lamina propria of the gastrointestinal tract where it plays a critical role in the humoral response against gut infections [39]. Second, IgA is an important contributor to the regulation of helminth abundance and vital rates [40–45], where the degree of protection is strongly affected by the host-helminth system and its history of infection [46–48]. In this system, we found that IgA (but not IgG) follows the dynamics of the two helminths, suggesting a rapid response and a reasonable representation of the immune response that does exhibit some regulatory proprieties but no long-term protection [23–27]. Third and most important, we needed a variable with an antigenic-specific immune response that could allow the quantification of separate reactions, namely species-specific and cross-reacting signals and, as such, would capture the essence of symmetric/asymmetric interactions, while avoiding additional complexities of adding extra variables and unnecessary assumptions. Our selection was also for a variable that could be easily quantified from large field datasets. Moreover, antibodies are tractable model tools since we can select among different isotypes based on the parasite-specific relationship of interest.

Species-specific serum IgA was originally quantified using excretory/secretory products of adult parasites, as a source of antigen, and the enzyme-linked immunosorbent assay (ELISA) [29]. Tests of *in vitro* antibody performance and competitive abilities between the two helminths indicated good selective abilities to discriminate between the two helminths when they co-occur [29]. Despite these in vitro analyses, we did not exclude the possibility of antibody cross-reactivity, especially at high intensities of infection. Finally, weather variables on mean air temperature and relative humidity were collected daily from the nearby Hutton Institute (UK).

### Model Framework

We used a deterministic, age-structured, immuno-epidemiological model with climatic forcing, previously selected among different candidate models for this rabbit-helminth system [30,31]. The model was then expanded to analyse hosts with single and dual infections and to explicitly include the contribution of species-specific and cross-reacting IgA responses. The demography of the host population was also included. We assumed that the within-host helminth dynamic can be captured as:

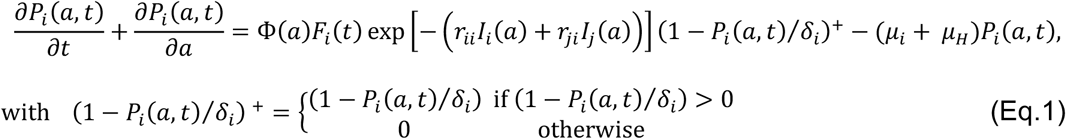

where: the intensity of infection (IOI) of the focal helminth *P_i_* (*i* = *T* for *T. retortaeformis* or *i* = *G* for *G. strigosum)*, changes with host age, *a* in days, and time, *t*. The intensity of the helminth-specific IgA response, *I_i_*, changes with host age, *a*; 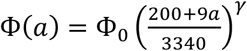 is an allometric function describing the age-dependent host feeding rate, with Φ_0_ and *γ* being suitable constants for each helminth species [30]. Given the similar IgA responses to infective third stage larvae (L3) and adults [24,25], and to avoid model redundancy, we focused on adults, assuming that there is no time delay from the ingestion of L3 to parasite maturation. The *r* parameters account for the effect of the immune response on the parasite intensity of infection, specifically, *r_ii_I*(*a*) is the age-dependent and species-specific IgA response that is stimulated by, and targets, parasite *i*, while *r_ji_I*(*a*) represents the age-dependent and species-specific IgA stimulated by parasite *j* (*j*≠*i*) that targets parasite *i* in dual-infected rabbits. *r_ii_* and *r_ji_* regulate the intensity of the stimulus triggered by the parasite; setting *r_ji_*=0 (*j*≠*i*) accounts for single infections. IgA is modelled to impact within-host parasite establishment and survival. Since the two helminths colonize different niches we did not consider direct inter-specific competition although we included the non-negative term (1 - *p_i_*(*a, t*)/*δ_i_*)^+^, which quantifies the intra-specific intensity-dependent effect on helminth establishment, with *δ_i_* representing the carrying capacity of helminth species *i. μ_i_* is the species-specific natural mortality rate of established parasites while *μ_H_* is the host natural mortality rate (*μ_H_* = 0.0069 days^-1^) [38]; the total mortality rate of parasites *μ* includes both *μ_i_* and *μ_H_*.

*F_i_*(*t*) is the risk of infection (RI) of the host by parasite *i* and quantifies the density of free-living helminths available for infection on the herbage at time *t. F_i_*(*t*) is driven by the intensity of infection in the host, *P_i_*, and the linear effects of air temperature, *τ*(*t*), and relative humidity, *H*(*t*), on the mortality of free-living stages, *μ_Fi_*(*τ*(*t*),*H*(*t*)) [31] as:

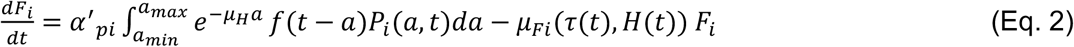

with *α’_pi_* = *α_pi_s_i_R*. The quantity *α’_pi_* describes the rate at which eggs are shed by an infected host and includes the total annual recruitment of rabbits, *R*, into the host population, the total number of eggs shed by an adult parasite per unit of time, *s_i_*, independent of host age, and the survival of free-living stages, *α_pi_*. In the model, eggs hatch directly into infective stages to help reduce model complexity while retaining the fundamental biological characteristics of the system and the emphasis on the within-host processes, and yet permitting an estimate of parasite fitness. The relative impact of weather on the loss of free-living stages is set as: *μ_Fi_*(*τ*(*t*), *H*(*t*)) = *α*_0*i*_—*α*_1*τi*_*τ*(*t*)—*α*_1*hi*_*H*(*t*), where *α*_0*i*_ represents the baseline natural parasite mortality rate, while *α*_1*τi*_ and *α*_1*hi*_ depict changes in the mortality rates driven by temperature and humidity, respectively [31]. The proportional change of *F_i_*(*t*) with the host intensity of infection is described by the term 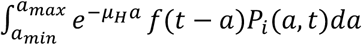, where *a_max_* = 283 days is the maximum age of an individual recorded in the data and *a_min_* = 30 days, which is approximately the age when naïve rabbits switch from milk to herbage and are exposed to infective stages. The first age class of naïve rabbits is initiated by assigning a null intensity of both infection and IgA. *F_i_*(*t*) explicitly depends on the age structure of the host population and thus, accounts for host reproduction, *f*(*t*), the relative number of births at time *t*, and host survival, *e^-μ_H_a^*, which represents the probability that rabbits are still alive at age *a*. We modelled *f*(*t*) as a beta probability density function calibrated against the fraction of 2-months old rabbits (SI-2.1). We assumed that infections with the two helminths occur through simultaneous ingestion, which is commonly expected in natural conditions; eggs are also shed simultaneously [29]. Table 1 summarizes the model variables and parameters.

**Table 1.** Variable and parameter definitions, and their units for model equations 1 and 2. The symbols *a* and *t* represent age and time in days.

| Variable/<br>parameter | Definition | Unit |
| --- | --- | --- |
| $P(a,t)$ | Intensity of infection | parasites * rabbit<sup>-1</sup> |
| $F(t)$ | Risk of infection | parasites * (grass unit)<sup>-1</sup> |
| $\Phi(a)$ | Age-dependent host feeding rate | grass unit * (day * rabbit)<sup>-1</sup> |
| $I(a)$ | Age-dependent host immune response | - |
| $\gamma$ | Allometric exponent of the host feeding rate | - |
| $\Phi_0$ | Host feeding rate for reference body mass of 3340g | grass unit * (day * rabbit)<sup>-1</sup> |
| $\mu_i$ | Natural mortality rate of parasite $i$ within the host | days<sup>-1</sup> |
| $\mu_H$ | Host natural mortality rate | days<sup>-1</sup> |
| $\delta$ | Parasite carrying-capacity | (parasites * rabbit<sup>-1</sup>)<sup>-1</sup> |
| $\mu_F$ | Weather-dependent mortality rate of free-living parasites | days<sup>-1</sup> |
| $\alpha_0$ | Parasite baseline mortality rate in the environment associated with worm features | days<sup>-1</sup> |
| $\alpha_{1\tau}$ | Effect of temperature on mortality rate of free-living parasites | days<sup>-1</sup> * °C<sup>-1</sup> |
| $\alpha_{1h}$ | Effect of humidity on mortality rate of free-living parasites | days<sup>-1</sup> * units of relative humidity<sup>-1</sup> |
| $\tau(t)$ | Mean daily air temperature from min and max records | °C |
| $H(t)$ | Daily relative air humidity from dry-web bulb temperature, atmospheric pressure at 101.3kPa | % |
| $\alpha'_p$ | Parasite shedding rate * host total recruitment | rabbit * (grass unit * days)<sup>-1</sup> |
| $r_{ii}$ | Intensity of immune response to parasite $i$ triggered by $i$ | - |
| $r_{ji}$ | Intensity of immune response to parasite $i$ triggered by $j$ | - |
| $k$ | Parasite aggregation from negative binomial distribution | - |

As the framework (Eqs. 1 and 2) is rather complex, and to identify the model that represents the best compromise between complexity and data availability, we focused on population A dataset to quantify the functional relationships and fundamental mechanisms of helminth dynamics, and used population B for the details on the host immune response. The first nontrivial problem we faced was fitting the model to data from population A with all the observed combinations of helminth abundances in dual-infected rabbits, including changes in the proportion of hosts with single and dual infections over time. To reduce this computational difficulty, the rabbit population was grouped in four subsets of infection data and the model was independently fitted to individual rabbits from each of these subsets: i)- rabbits with *T. retortaeformis* and free of *G. strigosum*, ii)- rabbits with *G. strigosum* and free of *T. retortaeformis*; iii)- rabbits with both parasites but fitting to *T. retortaeformis* data only and iv)- rabbits with both parasites but fitting to *G. strigosum* data only. Animals free of both helminths at the time of sampling, i.e. not currently infected, were included in every subset to provide the naïve condition, once dubious cases were identified and removed.

A similar approach was used for assigning the IgA values to rabbits from population A. The IgA data from population B were initially grouped into the four subsets and, for each one, the continuous age-related immune values *I_i_*(a) and *I_j_*(a) were obtained by interpolating a 4^th^ order polynomial function to the relationship between mean IgA and host age, weighted by sample size (figure S2*a* and *b*). Different smoothing functions were examined and the 4^th^ order polynomial fitted well our data. Rabbits from population A were then assigned an interpolated IgA value according to their type of infection and age.

The second non-trivial problem was related to model selection [49] and the possible overfitting when using a model with many free parameters. We tested five hypothesis-driven models (three for single infections) that represented different mechanisms of parasite regulation: i)- no constraints and parasites are affected by birth-death processes, ii)- immune-mediated constraints through IgA responses, iii)- intra-specific intensity-dependent constraints for host’s resources and, for the dual-infected hosts, we also examined iv)- cross-immunity via IgA responses (table 2). The complexity of the framework was explored by considering models that included these mechanisms in different combinations. Model selection and parameter calibration were performed independently for each of the four subsets using individual data from population A.

**Table 2.** Tested hypotheses and related mechanisms for single and dual infections of both helminths with the corresponding model complexity *p*, indicated by the number of parameters to be calibrated for each data subset.

| Models | Single infection | Dual infection | Parameter constraints | $p$ |
| --- | --- | --- | --- | --- |
| M0 | Parasite Birth-Death only | Parasite Birth-Death only | $r_{ji} = 0$ for any $j$ and $i$;<br>$\delta_i = \infty$ | 8 |
| M1 | Birth-Death + Specific Imm. | Birth-Death + Specific Imm. | $r_{ji} = 0$ for $j \neq i$; $\delta_i = \infty$ | 9 |
| M2 | Birth-Death + Specific Imm.<br>+ Intensity Depend. | Birth-Death + Specific Imm.<br>+ Intensity Depend. | $r_{ji} = 0$ for $j \neq i$ | 10 |
| M3 | - | Birth-Death + Specific Imm. +<br>Cross Imm. | $\delta_i = \infty$ | 10 |
| M4 | - | Birth-Death + Specific Imm. +<br>Cross Imm. + Intensity Depend. | - | 11 |

To increase model adherence to observed data the shape parameter *γ*, pertaining the rabbit’s feeding rate, Φ(*a*), was initially tuned by fitting the most complex model (M2 for single and M4 for dual infections) to the annual mean intensities of infection by host age, simultaneously in single and dual infections of each helminth species. We then used this value for all the models, including the less complex ones. During model fitting, *γ* was kept fixed while all the other parameters were recalibrated to the individual data of each of the four subsets. The best-performing model formulation was selected using the Akaike Information Criterion [49] and the lowest AIC. To allow the system to reach regime conditions during calibration, the model was simulated for an initial warm-up of 23-years that was then removed, full details of this procedure are reported elsewhere [31].

The numerical integration of the model was achieved using MATLAB^®^ *ode45* solver function based on an explicit Runge-Kutta solution with adaptive time step size [50]. For each of the four subsets of data, we compared the empirical helminth intensity in every rabbit to the expected intensity of infection by time and age of every host provided by the model output, assuming that the intensity of infection is distributed as a negative binomial [28] with age- and time-dependent mean *P*(*a,t*) and aggregation parameter *k*, calibrated together with the model parameters [28]. The parameters combination that maximized the likelihood function was selected using a non-linear solver based on the Nelder-Mead simplex algorithm [51].

Finally, to provide a statistical measurement of the differences between single and dual infected rabbits, the model simulated quantities of interest, namely, intensity of infection, abundance of eggs shed, risk of infection (i.e. viable free-living stages) and antibody response, were compared using Generalized Linear Models (GLM, with normal or negative binomial distribution of errors). Type of infection (single or dual) was entered as categorical variable, while host age was included as a continuous variable. The additive effects and two-way interactions of the explanatory variables were examined. For consistency, the same analysis was repeated using the empirical data (SI-1.2).

## Results

Here we present simulations from the best selected model and parameter calibration on population A, the focus of our study. All the statistical results are reported in SI-3.

### Epidemiology of Single and Dual infections

The model that best described the four subsets of infection data comprised the species-specific IgA response and the intra-specific intensity-dependence for single infections, dual infections also included IgA cross-reaction between the two helminths (table 2, SI-3.1). Individual-based simulations captured the average trends of infection by host age (figure 2), but less so the large intra-annual variation observed in the empirical data (figure S6). For both helminths the simulated intensities by host age were significantly higher in rabbits with dual than single infections (table S3). *T. retortaeformis* rapidly accumulated with host age and maintains high intensities in older rabbits (figure 2*a*), while accumulation of *G. strigosum* intensities was slower (figure 2*b*). Consistent with these model results, the empirical intensities of infection by host age were significantly higher in rabbits with dual than single infection, for both helminths (figure 2, table S3).

### Immune response and Immune-mediated parasite facilitation

The estimation of immunological parameters yields information on the immunological mechanism that could generate the epidemiological patterns observed. Simulations indicate that the stimulus to develop a specific IgA response to *T. retortaeformis* is stronger in rabbits with single than dual infections (*r_TT_*= 0.734 vs 0.338, table 3). In this latter group, the cross-reaction stimulus for a specific IgA response against *G. strigosum* that also attacks *T. retortaeformis* is essentially null (*r_GT_*= 0.00016). The projected relationships between intensity of infection and IgA stimuli indicate that a proportional increase in the specific IgA stimulus (*r_TT_*) will reduces *T. retortaeformis* intensities, while the stimulus for an IgA response to *G. strigosum* that cross-reacts with *T. retortaeformis* (*r_GT_*) remains extremely weak and will have no impact on this helminth (figure 3*a*). When we consider the estimated species-specific antibody response against *T. retortaeformis, r_TT_I*, values are significantly lower in dual than single infection, and for rabbits of all age groups (figure 4*a*, table S4). These results suggest that the higher *T. retortaeformis* intensities in dual-infected hosts are caused by the weakening of the stimulus to a specific IgA response that is diverted against *G. strigosum* (also see below) and a negligible immune-mediated interference by this latter helminth in the form of a weak cross-reaction, *r_GT_I*. We note that, while empirical data only quantify specific IgA’s, the model allows us to quantify both the species-specific (*r_ii_I_i_*) and the cross-reactive (*r_ji_I_j_*) relative strength of IgA, namely the effect of an increase (or decrease) of IgA on the intensity of infection.

**Figure 3.**
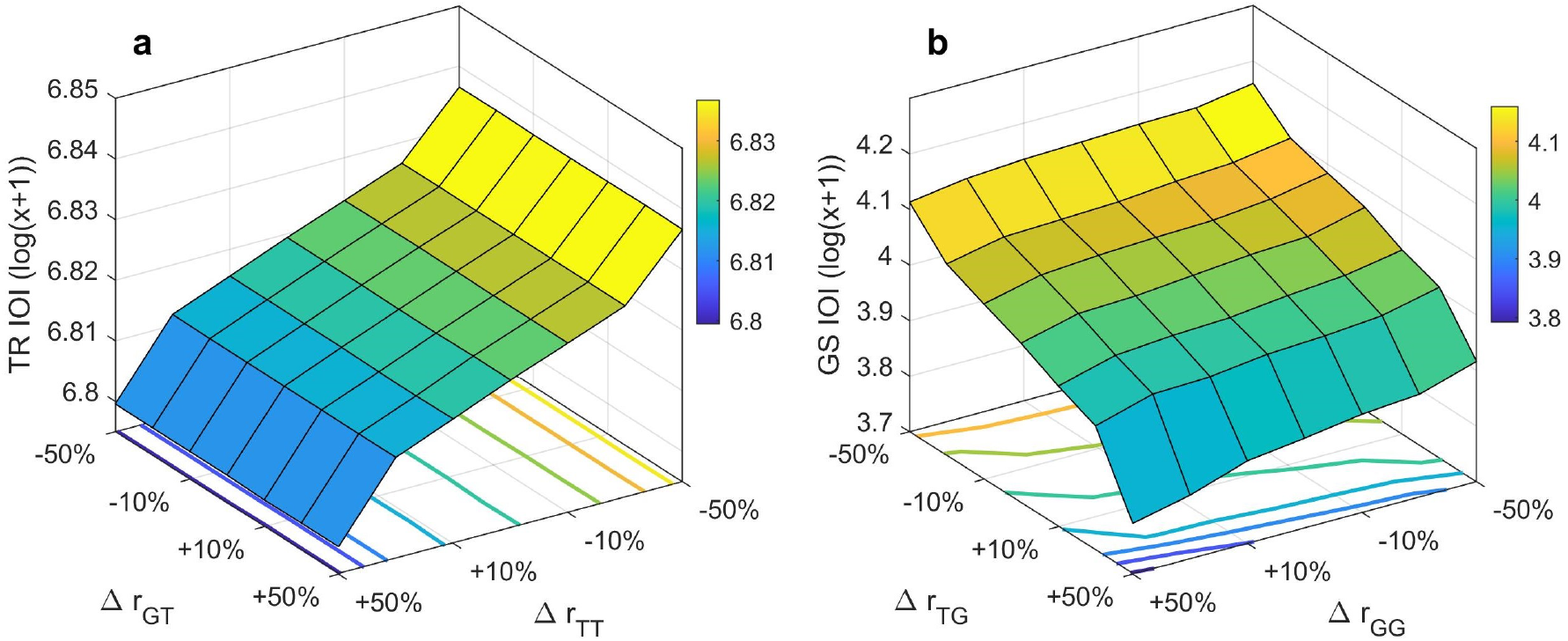
Three-way relationships among model-predicted intensity of infection (IOI) and relative changes in species-specific (Δ*r_TT_* or Δ*r_GG_*) and cross-reacting (Δ*r_TG_* or Δ*r_GT_*) stimuli to IgA production for *T. retortaeformis* (TR, a) and *G. strigosum* (GS, b). Predictions are from the selected best model for dual infections (M4) fitted on population A. The bivariate incremental variation (% of increase or decrease) in the immune parameters, *r_ii_* or *r_ji_*, is relative to the baseline estimated values reported in table 3. The IOIs (heat surface) are obtained via sensitivity analysis where the immune parameters are incrementally changed one at a time, with respect to their estimated values, and simulations are run while holding all the other parameters constant. The contour lines represent the predicted IOI values from the heat map at specific *r_ii_* and *r_ji_* values.

**Figure 4.**
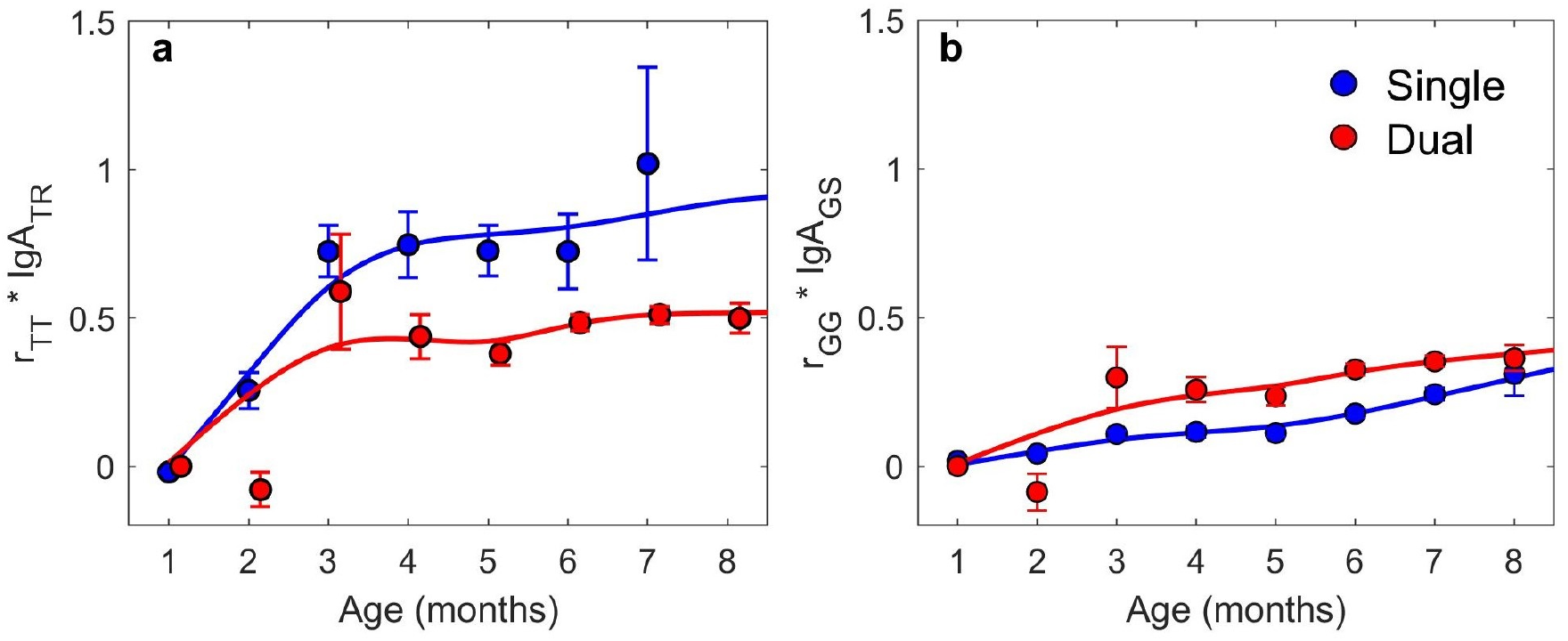
Relationship between the contribution of species-specific IgA response, r*_ii_*IgA, by host age for *T. retortaeformis* (a) and *G. strigosum* (b) in single- (blue) and dual- (red) infected hosts from population A as inferred from the model. The simulated r*_ii_*IgA values (mean and S.E.) and the 4^th^ order polynomial curves, weighted by sample size, are reported. Small S.E. are masked by the circles.

**Table 3.** Parameter values estimated from the best-fitted model for *T. retortaeformis* (TR) and *G. strigosum* (GS) in single- (SI) and dual- (DU) infected hosts. Parameter definitions are reported in table 1. *F*(0) refers to the initial condition of the warm-up period of model simulations while *γ* is included as a constant value. AIC= Akaike Information Criterion value, LL= log-likelihood, RMSD= Root Mean Square Deviation.

| Parameters | TR-SI | TR-DU | GS-SI | GS-DU |
| --- | --- | --- | --- | --- |
| $\gamma$ | 3.09 | 3.09 | 5.73 | 5.73 |
| $\Phi_0$ | 1.70 | 0.4738 | 1.84 | 0.430 |
| $\mu_i$ | 0.0040 | 0.00219 | 0.212 | 0.0161 |
| $\delta$ | 651.6 | 789.74 | 78.71 | 83.45 |
| $F(0)$ | 76.42 | 6.28 | 61.30 | 3.28 |
| $\alpha_0$ | 7.39 | 3.60 | 0.328 | 0.194 |
| $\alpha_{1\tau}$ | 0.445 | 0.207 | 0.0022 | 0.00019 |
| $\alpha_{1h}$ | -0.0003 | 0.00254 | 0.00375 | 0.00236 |
| $\alpha_p'$ | 1.86 | 3.322 | 0.129 | 0.1323 |
| $k$ | 0.220 | 0.244 | 0.146 | 0.423 |
| $r_{ii}$ | 0.734 | 0.338 | 0.1206 | 0.208 |
| $r_{ji}$ | - | 0.00016 | - | 0.804 |
| Parameters | 10 | 11 | 10 | 11 |
| AIC | 17,074 | 19,204 | 3,052 | 10,878 |
| LL | 8,624 | 9,591 | 1,516 | 5,428 |
| RMSD | 576 | 999 | 49.2 | 73.4 |

The intra-specific carrying capacity, *δ*, is higher in rabbits with dual than single infections (789.74 vs 651.6, respectively, table 3) supporting the lower intensity-dependent control in this latter group.

For *G. strigosum*, simulations showed that the stimulus to develop a specific IgA response is low (table 3), and lower in rabbits with single (*r_GG_*= 0.121) than dual infection (*r_GG_*= 0.208). The stimulus for an IgA response specific to *T. retortaeformis* that cross-reacts against *G. strigosum* is four times higher (*r_TG_*= 0.804), implying interference from this helminth. Moreover, the investigation of how *G. strigosum* intensities will change in relation to changes in the antibody stimuli indicates that the simulated intensities will decline with a proportional increase of *r_TG_* and less so of *r_GG_* (figure 3*b*). Examination of the estimated antibody response, *r_GG_I*, by host age showed that values are significantly higher in rabbits with dual than single infections (figure 4*b*, table S4), although it is important to observe that these values remain consistently low. All together, these findings suggest that dual-infected rabbits should control *G. strigosum* more successfully than single-infected hosts. However, the generally low IgA response, even for the cross-reaction, could explain the significantly higher intensities observed in dual-infected hosts (table S3).

Previous studies proposed that intensity-dependence could be more important for the dynamics of this parasite [31]. Our simulations indicate that parasite carrying capacity is slightly higher in hosts with dual than single infections (*δ*= 83.45 and 78.71, respectively, table 3), suggesting the tendency for weaker restrictions of *G. strigosum* intensities in this latter group, as previously noted for *T. retortaeformis*.

Collectively, the weakening or fundamentally low IgA responses, and the resulting higher intensities of both helminths in dual infections, support the hypothesis of immune-mediated facilitation between the two species. These higher intensities are generated despite a clear uneven in the immune response against the two helminths.

### Parasite fitness and Risk of infection

Given the soil-transmitted nature of *T. retortaeformis* and *G. strigosum*, the density of viable free-living stages on the pasture represents the risk of infection for hosts exposed to these stages, and can be considered a proxy for onward transmission and parasite fitness. The density of free-living stages is the sum of the abundance of eggs shed by every host and their mortality rate. We used simulation results to examine whether parasite immune-mediated facilitation has symmetrical or asymmetrical consequences for parasite fitness. The estimated abundance of eggs shed on the pasture is significantly higher in rabbits with dual than single infections, for both helminths (table S5). Eggs shedding peaks in young and a decrease in older animals for *T. retortaeformis* while there is a constant increase with host age for *G. strigosum* (figure S7*a,b*).

On the herbage, the population of free-living stages is affected by environmental factors (biotic and abiotic) with higher mortalities for *T. retortaeformis* than *G. strigosum* (table 3). Throughout the study period, the mortality rate of free-living stages produced by rabbits with single and dual infections is, respectively: *μ_FT_*= 3.52 and 1.58 day^-1^ for *T. retortaeformis* and *μ_FG_* = 0.009 and 0.003 days^-1^ for *G. strigosum*. Part of this loss is driven by weather: temperature has a strong impact on *T. retortaeformis* (*α*_1*τ*_ = 0.445 and 0.207 for single and dual infections, respectively) while humidity is more relevant to *G. strigosum* (*α*_1*H*_ = 0.004 and 0.002). The ecological differences between the two helminths are also clear from the baseline natural mortality rate of free-living stages, *α*_0_, which is a component of *μ_Fi_*. Mortality is higher for *T. retortaeformis* (*α*_0_ = 7.39 and 3.60 for single and dual infections, respectively) than for *G. strigosum* (*α*_0_ = 0.328 and 0.194), indicating that larvae are on the pasture for a shorter period of time for the former helminth. More important, simulations suggest that under analogous environmental conditions, free-living stages from both helminths experience lower natural and climate-driven mortality, and thus remain available for onward transmission for longer, when derived from rabbits with dual infections (compare *α*_0_ and *μ_Fi_* from single and dual infections).

There is strong seasonality in the accumulation of free-living stages on the herbage, and hence the risk of infection (figure 5). *T. retortaeformis* accumulation is at the highest in August primarily through the shedding of young and newly infected rabbits and drops in June, coinciding with the peak of newly born rabbits (figure S3). *G. strigosum* shows the highest densities around March, when the population is composed predominantly of adults and the lowest around July, especially for dual infected hosts, with the arrival of newborn hosts. In both cases, rabbits dual-infected generate significantly more viable free-living stages throughout the year, than single-infected rabbits (table S6). Therefore, while differences between the two helminths are expected because of their ecological characteristics, both *T. retortaeformis* and *G. strigosum* from dual-infected rabbits exhibit significantly higher fitness, including higher intensities of infection, than helminths from single infections, supporting the hypothesis of symmetrical immune-mediated facilitation between the two parasites.

**Figure 5.**
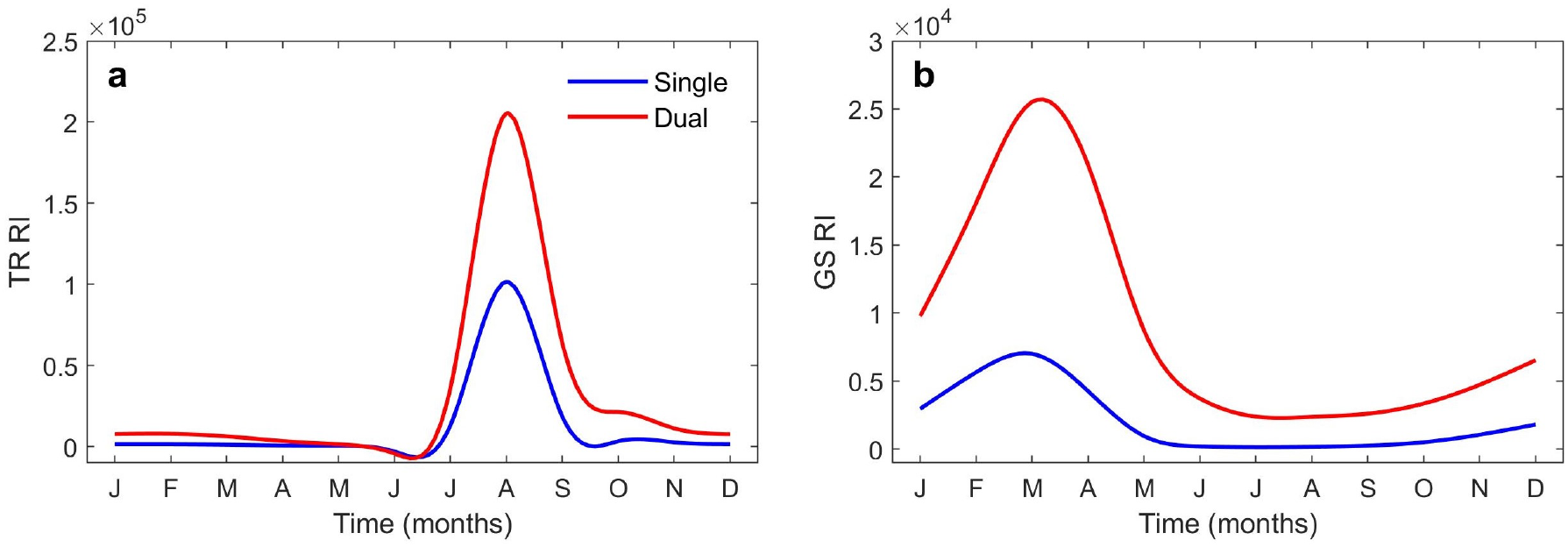
Estimated seasonality of the mean risk of infection (RI) by sampling month for *T. retortaeformis* (TR, a) and *G. strigosum* (GS, b) in single- (blue) and dual-infected (red) hosts from population A.

## Discussion

We applied an immuno-epidemiological model to empirical data of two helminths from a population of rabbits and the findings support the hypothesis that immune-mediated facilitation can explain *T. retortaeformis* and *G. strigosum* higher intensities in rabbits with dual compared to single infections. Dual infections are facilitated by weakened species-specific IgA responses and unequal IgA cross reactions. Our multi-scale approach also suggests a symmetrical immune-mediated facilitation where weak IgA stimuli contribute to greater number of eggs shed and higher survival of the free-living stages, and thus higher transmission, when compared to parasites derived from hosts with single infections. Given that rabbits with dual infections represent the large majority of the sampled population (figure S4), the dynamics of the two helminths is primarily driven by this group of hosts.

Cross-immunity, where protection to one species provides some defence against a second species, has been proposed in a variety of systems [e.g. 1-3,6,52]. Our study offers novel insights by showing that, although some immune-mediated interference between two helminths is likely to occur, IgA cross-reacts disproportionately and has small impact on parasite intensity, it is essentially null against *T. retortaeformis* and has a low effect against *G. strigosum*. This pattern is consistent with theoretical work showing that two parasites will increase abundance and persist in the host if immune-mediated facilitation is not too strong to destabilize the system [19]. The evidence in animal systems of positive interactions between parasite species [1,2,53], suggests some level of immune-mediated facilitation, irrespective of the specific immune mechanism involved. Examples of facilitation through host immunity have been frequently described for HIV associated co-infections in humans, such as HIV-malaria or HIV-Tb [58,59]. Positive interactions between micro- and macro-parasites, mediated by trade-offs in the immune functions and responses, are expected to benefit one or both parasites [11,13]. For helminth co-infections, synergistic effects could emerge throughout diverse processes; for example, the allocation of IgE against helminth species could help explain the positive correlation between *Ascaris lumbricoides* and both *Trichuris trichiura* and hookworm infections in humans (15, 54-56). Similarly, the co-circulation of *Trichostrongylus* spp., *Haemonchus contortus* and *Teladorsagia circumcincta* in domestic and wild animal populations is facilitated by their immuno-modulatory properties [57] and could be complemented by asymmetric immune reactions.

We can expect that one of the consequences of immune-mediated facilitation will be greater parasite fitness. We found that rabbits with dual-infections contribute to a larger number of free-living stages that are available for onward transmission, than did rabbits with single infection. Our model did not explicitly quantify the relationships between IgA and parasite size or fecundity, where both fecundity and shedding are directly related to parasite body length [23,29], but coupled shedding to infection intensity, which is then modulated by IgA. Some of these relationships were previously examined and showed a negative relationship between IgA and *G. strigosum* body length in single infections and between IgA and *T. retortaeformis* body length or abundance in co-infections [23]. A combination of positive and negative relationships was also found between the vital rates of both helminths and type 1, type 2 and T-regulatory immune variables [23], confirming the complex interactions between parasite demography and the host immune response. Indeed, the immune response to an infection is the results of a large number of functions and factors, each of which has distinct roles and degree of specificity. Our model framework describes a small constituent of this immune network, and gratifyingly captured the effects, while other immune processes could also have potentially contributed to the weaker net response of dual infected rabbits. More broadly, our results are consistent with studies from other systems on the importance of IgA to helminth growth, fecundity and shedding [60–63].

In addition to the diverse tolerance of the two helminths to weather [33–35], the higher natural mortality of *T. retortaeformis* on the pasture is probably associated with the faster life history of this parasite, namely, faster egg hatching rate [34] and faster within-host maturation [64], when compared to *G. strigosum* [65]. Crucially, free-living stages derived from rabbits with dual infections exhibited longer survival leading to higher probability of onwards transmission. Laboratory studies showed that fewer antibodies bind to eggs of *T. retortaeformis* than *G. strigosum*, and egg volume decreases for the first and increases for the second during a single-dose infection experiment [66]. While there was no clear evidence that antibodies altered egg size or hatchability, the impact could be relevant in natural settings, when both the host and the helminths are under environmental constraints. For instance, the low IgA specific response to *T. retortaeformis* in hosts with dual-infections could lead to eggs of higher quality, such as larger size or more tolerant to thermal changes, by females in better conditions. For *G. strigosum* the mechanism is less clear but could also be related to improved female conditions and enhanced egg quality with better tolerance to moisture variation.

Our data-informed modelling approach contributes to advances in the understanding of immune-mediated facilitation on infection and transmission. The mechanistic understanding of these processes, and the contribution to parasite dynamics and fitness, is still limited and in great need of empirical evidence. There is also a need to improve the realism of modelling multiple infections, particularly for macro-parasites. The proposed framework can be adapted to test alternative and more complex immune-mediated formulations beyond the species examined in this study. *T. retortaeformis* and *G. strigosum* share similarities with other gastrointestinal helminths of animals, including human parasites, and our findings have relevance across a broad range of ecological settings. The fundamental challenge is to identify the key variables that can clarify the mechanisms of regulation across scales and influences parasite fitness. Gathering this information can be daunting but is a prerequisite for laying the foundation of a better understanding of the ecological role of co-infection in disease spread and persistence, which is essential for developing control measures tailored on these groups of hosts.

## Acknowledgments

This study was supported by the National Science Foundation (DEB-1145697). FD was partially sponsored by The Ermenegildo Zegna Founder’s Scholarship. We are grateful to Tricia Brockman for creating figure S1.

## Ethic statement

We used rabbit data already available from previous studies where sampling was performed according to field procedures approved by the Institutional Animal Care and Use Committee of The Pennsylvania State University (IACUC # 26383 and 34489). All animal work adhered to the guidelines laid out in the Guide for the Care and Use of Laboratory Animals. 8th ed. National Research Council of the National Academies. National Academies Press Washington DC.

## References

1. Cox, F.E.G. 2001. Concomitant infections, parasites and immune responses. Parasitology 122, S23–S38.

2. Christensen, N.Ø., P. Nansen, B.O. Fagbemi and J. Monrad. 1987. Heterologous antagonistic and synergistic interactions between helminths and between helminths and protozoans in concurrent experimental infection of mammalian hosts. Parasit. Res. 73, 387–410.

3. Lee, T.D.G., R. K. Grencis and D. Wakelin. 1982. Specific cross-immunity between *Trichinella spiralis* and *Trichuris muris:* immunization with heterologous infections and antigens and transfer of immunity with heterologous immune mesenteric lymph node cells. Parasitology 84, 381–389.

4. Gupta, S., J. Swinton, and R.M. Anderson. 1994. Theoretical studies of the effects of heterogeneity in the parasite population on the transmission dynamics of malaria. Proc. R. Soc. B. 256, 231–238.

5. Råberg, L., J.C. De Roode, A.S. Bell, P. Stamou, D. Gray, and A.F. Read. 2006. The role of immune-mediated apparent competition in genetically diverse malaria infections. Am. Nat. 168, 41–53.

6. Bhattacharyya, S., P.H. Gesteland, K. Korgenski, O.N. Bjørnstad, and F.R. Adler. 2015. Cross-immunity between strains explains the dynamical pattern of paramyxoviruses. PNAS 112, 13396–13400.

7. Zélé, F., S. Magalhães, S. Kéfi, and A.B. Duncan. 2018. Ecology and evolution of facilitation among symbionts. Nat. Com. 9, 4869.

8. McSorley, H.J. and Maizels, R.M., 2012. Helminth infections and host immune regulation. Clin. Microbiol. Rev. 25, 585–608.

9. Behnke, J.M., Wahid, F.N., Grencis, R.K., Else, K.J., Ben-Smith, A.W. and Goyal, P.K., 1993. Immunological relationships during primary infection with *Heligmosomoides polygyrus (Nematospiroides dubius):* downregulation of specific cytokine secretion (IL-9 and IL-10) correlates with poor mastocytosis and chronic survival of adult worms. Parasi. Immunol. 15, 415–421.

10. Genta, R.M. and Walzer, P.D., 1989. Strongyloidiasis. In Parasitic Infections in the Compromised Host (ed. Walzer, P. D. & Genta, R. M.), pp. 463–525. New York: Marcel Dekker.

11. Cattadori, I.M., R. Albert, and B. Boag. 2007. Variation in host susceptibility and infectiousness generated by co-infection: the *myxoma-Trichostrongylus retortaeformis* case in wild rabbits. J.R.S. Interface 4, 831–840.

12. Bickle, Q. D., Solum, J., & Helmby, H. (2008). Chronic intestinal nematode infection exacerbates experimental *Schistosoma mansoni* infection. Infect. Immun. 76, 5802–5809.

13. Graham A. 2008. Ecological rules governing helminth-microparasite coinfection. PNAS 105, 566–570.

14. Onah D.N. and Wakelin D., 1999. Trypanosome-induced suppression of responses to *Trichinella spiralis* in vaccinated mice. Int. J. Parasit. 29, 1017–1026.

15. Fleming, F.M., S. Brooker, S.M. Geiger, I.R. Caldas, R. Correa-Oliveira, P.J. Hotez, and J.M. Bethony. 2006. Synergistic associations between hookworm and other helminth species in a rural community in Brazil. Trop. Med. Intern. Health 11, 56–64.

16. Diedrich C.R. and J.L. Flynn 2011. HIV-1/mycobacterium tuberculosis coinfection immunology: how does HIV-1 exacerbate tuberculosis? Infect. Immun. 79, 1407–1417.

17. Shi, K., Li, H., Guo, X., Ge, X., Jia, H., Zheng, S. and Yang, H., 2008. Changes in peripheral blood leukocyte subpopulations in piglets co-infected experimentally with porcine reproductive and respiratory syndrome virus and porcine circovirus type 2. Vet. Microbiol. 129, 367–377.

18. Soares M.P., Teixeira L. and Moita L.F., 2017. Disease tolerance and immunity in host protection against infection. Nat. Rev. Immunol. 17, 83–96.

19. Eswarappa, S.M., S. Estrela and S.P. Brown. 2012. Within-host dynamics of multi-species infections: facilitation, competition and virulence. PLoS One 7: 38730.

20. Kamiya, T., N. Mideo, and S. Alizon. 2018. Coevolution of virulence and immunosuppression in multiple infections. J. Evol. Biol. 31, 995–1005.

21. Tillmann, H.L., Heiken, H., Knapik-Botor, A., Heringlake, S., Ockenga, J., Wilber, J.C., Goergen, B., Detmer, J., McMorrow, M., Stoll, M. and Schmidt, R.E., 2001. Infection with GB virus C and reduced mortality among HIV-infected patients. New Engl. J. Med., 345, 715–724.

22. Cattadori, I.M., B. Boag, O.N. Bjørnstad, S. Cornell, and P.J. Hudson. 2005. Peak shift and epidemiology in a seasonal host-nematode system. Proc. R. Soc. B. 272, 1163–1169.

23. Cattadori I.M., A.K. Pathak and M.J. Ferrari. 2019. External Disturbances Impact Helminth-Host Interactions by affecting Dynamics of Infection, Parasite Traits and Host Immune Responses. Ecol. Evol. 9, 13495–13505.

24. Murphy, L., N. Nalpas, M. Stear, and I.M. Cattadori. 2011. Explaining patterns of infection in free living populations using laboratory immune experiments. Paras. Immunol. 33, 287–302.

25. Murphy, L., A.K. Pathak, and I.M. Cattadori. 2013. A co-infection with two gastrointestinal nematodes alters host immune responses and only partially parasite dynamics. Paras.Immunol. 35, 421–432.

26. Vanalli C., L. Mari, R. Righetto, R. Casagrandi, M. Gatto, and I.M. Cattadori. 2020 Within-host mechanisms of immune regulation explain the contrasting dynamics of two helminth species in both single and dual infections. Plos Comp. Biol. 16, p.e1008438.

27. Thakar, J., A.K. Pathak, L. Murphy, R. Albert, and I.M. Cattadori. 2012. Network model of immune responses reveals key effectors to single and co-infection dynamics by a respiratory bacterium and a gastrointestinal helminth. PLoS Comp. Biol. 8, p.e1002345.

28. Cattadori, I.M., B. Boag, and P.J. Hudson. 2008. Parasite coinfection and interaction as drivers of host heterogeneity. Int. J. Parasit. 38, 371–380.

29. Cattadori, I.M., B.R. Wagner, L.A. Wodzinski, A.K. Pathak, A. Poole, and B. Boag. 2014. Infections do not predict shedding in co-infections with two helminths from a natural system. Ecology 95, 1684–1692.

30. Cornell, S.J., O.N. Bjornstad, I.M. Cattadori, B. Boag, and P.J. Hudson. 2008. Seasonality, cohort-dependence and the development of immunity in a natural host-nematode system. Proc. R. Soc. B 275, 511–518.

31. Mignatti, A., B. Boag, and I.M. Cattadori. 2016. Host immunity shapes the impact of climate changes on the dynamics of two parasite infections. PNAS 113, 2970–2975.

32. Cattadori, I.M., A. Sebastian, H. Hao, K. Katani, I. Albert, K.E. Eilertson, V. Kapur, A. Pathak and S. Mitchell. 2016. Impact of helminth infections and nutritional constraints on the small intestine microbiota. PLoS One 11, 0159770.

33. Boag, B., and R.J. Thomas. 1970. The development and survival of free-living stages of *Trichostrongylus colubriformis* and *Ostertagia circumcincta* on pasture. Res. Vet. Sci. 11, 380–381.

34. Hernandez, A.D., A. Poole, and I.M. Cattadori. 2013. Climate changes influence free-living stages of soil-transmitted parasites of European rabbits. Glob. Change Biol. 19, 1028–1042.

35. Gupta, S.P. 1961. The effects of temperature on the survival and development of the free-living stages of *Trichostrongylus retortaeformis* Zeder (Nematoda). Can. J. Zool. 391, 47–53.

36. Boag, B. 1985. The incidence of helminth parasites from the wild rabbit *Oryctolagus cuniculus* (L.) in Eastern Scotland. J. Helminth. 59, 61–69.

37. Pathak, A.K., C. Pelensky, B. Boag, and I.M. Cattadori. 2012. Immuno-epidemiology of chronic bacterial and helminth co-infections: observations from the field and evidence from the laboratory. Int. J. Parasit. 42, 647–655.

38. Smith, G.C., and R.C. Trout. 1994. Using Leslie matrices to determine wild rabbit population growth and the potential for control. J. Appl. Ecol. 31, 223–230.

39. Mestecky, J. and McGhee, J.R., 1987. Immunoglobulin A (IgA): molecular and cellular interactions involved in IgA biosynthesis and immune response. Adv. Immunol. 40, 153–245.

40. Sorobetea, D., M. Svensson-Frej, and R. Grencis. 2018. Immunity to gastrointestinal nematode infections. Muc. Immunol. 11, 304–315.

41. Ben-Smith, A., Wahid, F.N., Lammas, D.A. and Behnke, J.M., 1999. The relationship between circulating and intestinal *Heligmosomoides polygyrus-specific* IgG1 and IgA and resistance to primary infection. Paras. Immunol. 21, 383–396.

42. Van Knapen, F., J.H. Franchimont, A.R. Verdonk, J. Stumpf, and K. Undeutsch. 1982. Detection of specific immunoglobulins (IgG, IgM, IgA, IgE) and total IgE levels in human trichinosis by means of the enzyme-linked immunosorbent assay (ELISA). Am. J. Trop. Med. Hyg. 31, 973–976.

43. Clerc, M., Devevey, G., Fenton, A. and Pedersen, A.B., 2018. Antibodies and coinfection drive variation in nematode burdens in wild mice. Int. J. Parasit, 48, 785–792.

44. Escribano, C., Saravia, A., Costa, M., Castells, D., Ciappesoni, G., Riet-Correa, F. and Freire, T., 2019. Resistance to *Haemonchus contortus* in Corriedale sheep is associated to high parasite-specific IgA titer and a systemic Th2 immune response. Sci. Rep., 9, 1–10.

45. Inaba T, et al. 2003. Monoclonal IgA antibody-mediated expulsion of *Trichinella* from the intestine of mice. Parasitology 126, 591–598.

46. Esser-von Bieren, J., I. Mosconi, R. Guiet, A. Piersgilli, B. Volpe, F. Chen, W.C. Gause, A. Seitz, J.S. Verbeek, and N.L. Harris. 2013. Antibodies trap tissue migrating helminth larvae and prevent tissue damage by driving IL-4Rα-independent alternative differentiation of macrophages. PLoS Pathog. 9, e1003771.

47. McCoy, K.D., M. Stoel, R. Stettler, P. Merky, K. Fink, B.M. Senn, C. Schaer, J. Massacand, B. Odermatt, H.C. Oettgen, and R.M. Zinkernagel. 2008. Polyclonal and specific antibodies mediate protective immunity against enteric helminth infection. Cell Host Microb. 4, 362–373.

48. Roach, T.I.A., K.J. Else, D. Wakelin, D.J. McLaren, and R.K. Grencis. 1991. *Trichuris muris:* antigen recognition and transfer of immunity in mice by IgA monoclonal antibodies. Paras. Immunol. 13, 1–12.

49. Burnham, K.P., and D.R. Anderson. 2002. Model selection and multimodel inference: a practical information-theoretic approach. Springer, New York.

50. Dormand, J.R. and P.J. Prince. 1980. A family of embedded Runge-Kutta formulae. J. Comp. Appl. Math. 6, 19–26.

51. Lagarias, J.C., J.A. Reeds, M.H. Wright, and P.E. Wright. 1998. Convergence properties of the Nelder-Mead simplex method in low dimensions. SIAM J. Optim. 9, 112–147.

52. Johnson, P.T. and I.D. Buller. 2011. Parasite competition hidden by correlated coinfection: using surveys and experiments to understand parasite interactions. Ecology 92, 535–541.

53. Petney, T.N. and R.H. Andrews. 1998. Multiparasite communities in animals and humans: frequency, structure and pathogenic significance. Int. J. Parasit. 28, 377–393.

54. Booth, M., and D.A.P. Bundy. 1992. Comparative prevalences of *Ascaris lumbricoides, Trichuris trichiura* and hookworm infections and the prospects for combined control. Parasitology 105, 151–157.

55. Geiger, S.M., N.D.E. Alexander, R.T. Fujiwara, S. Brooker, B. Cundill, D.J. Diemert, R. Correa-Oliveira, and J.M. Bethony. 2011. *Necator americanus* and helminth co-infections: further down-modulation of hookworm-specific type 1 immune responses. PLoS Negl. Trop. Dis. 5, p.e1280.

56. Lepper, H.C., J.M. Prada, E.L. Davis, S.A. Gunawardena, and T.D. Hollingsworth. 2018. Complex interactions in soil-transmitted helminth co-infections from a cross-sectional study in Sri Lanka. Trans. R. Soc. Trop. Med. Hyg. 112, 397–404.

57. McNeilly, T.N., and A.J. Nisbet. 2014. Immune modulation by helminth parasites of ruminants: implications for vaccine development and host immune competence. Parasite 21.

58. Abu-Raddad, L.J., Patnaik, P. and Kublin, J.G., 2006. Dual infection with HIV and malaria fuels the spread of both diseases in sub-Saharan Africa. Science 314, 1603–1606.

59. Kwan, C.K. and J.D. Ernst. 2011. HIV and tuberculosis: a deadly human syndemic. Clin. Microb. Rev. 24, 351–376.

60. Stear, M.J., S.C. Bishop, M. Doligalska, J.L. Duncan, P.H. Holmes, J. Irvine, L. McCririe, Q.A. McKellar, E. Sinski, and M.A.X. Murray. 1995. Regulation of egg production, worm burden, worm length and worm fecundity by host responses in sheep infected with *Ostertagia circumcincta*. Parasit. Immunol. 17, 643–652.

61. Strain, S., S. Bishop, N. Henderson, A. Kerr, Q. Mackellar, S. Mitchell, and M. Stear. 2002. The genetic control of IgA against *Teladorsagia circumincta* and its association with parasite resistance in naturally infected sheep. Parasitology 124, 545–522.

62. Henderson, N.G., and M.C. Stear. 2006. Eosinophil and IgA responses in sheep infected with *Teladorsagia circumcincta*. Vet. Immunol. Immunopath. 112, 62–66.

63. McRae, K.M., M.J. Stear, B. Good, and O.M. Keane. 2015. The host immune response to gastrointestinal nematode infection in sheep. Paras. Immunol. 37, 605–613.

64. Audebert, F., H. Hoste, and M.C. Durette-Desset. 2002. Life cycle of *Trichostrongylus retortaeformis* in its natural host, the rabbit *(Oryctolagus cuniculus)*. J. Helminth. 76, 189–192.

65. Massoni, J., J. Cassone, M.C. Durette-Desset, and F. Audebert. 2011. Development of *Graphidium strigosum* (Nematoda, Haemonchidae) in its natural host, the rabbit *(Oryctolagus cuniculus)* and comparison with several Haemonchidae parasites of ruminants. Parasit. Res. 109, 25–36.

66. Lambert, K., A.K. Pathak. and I.M. Cattadori. 2005. Does host immunity influence hatchability of helminth eggs shed in the environment? J. Helminth. 89, 446–452.

