## Supplemental Information for "The Enemy of my Enemy is my Friend: Immune-Mediated Facilitation Contributes to Fitness of Co-Infecting Helminths"

SI-1 Empirical data page 2

1.1 Schematic of the helminth-rabbit system

1.2 Statistical analysis

1.3 Empirical data from population B

1.4 Structure of host population A and B

SI-2 Methods page 6

SI-2.1 Newborn recruitment in population A

SI-3 Results page 7

SI-3.1 Model selection on population A

SI-3.2 Annual intensities of infection of population A

SI-3.3 Intensities of infection and IgA response of population A

SI-3.4 Parasite shedding by population A

### SI-1 Empirical data

#### 1.1 Schematic of the helminth-rabbit system

The graphic of the dynamics of infection of the gastrointestinal helminths *T. retortaeformis* and *G. strigosum* in rabbits with single and dual infections, including key components of the dynamics, such as antibody mediated regulation and the role of weather on stages free-living on the herbage, are depicted in figure S1.

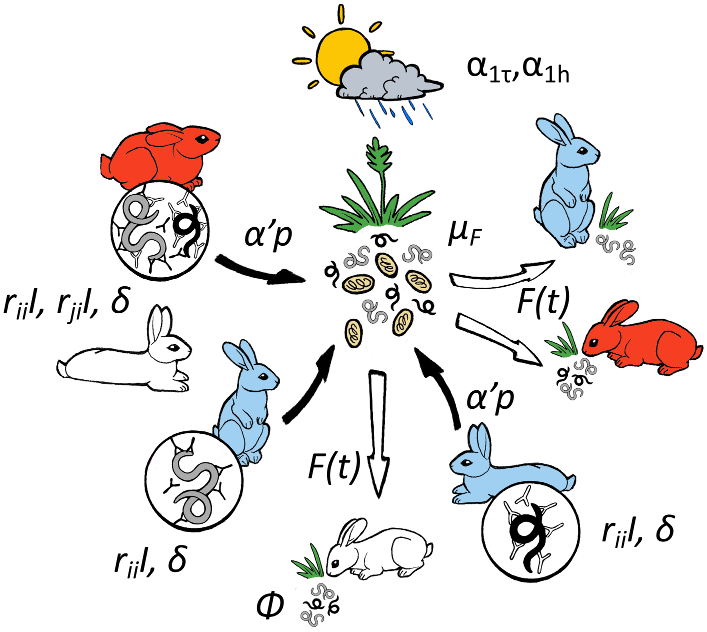


**Figure S1.** Schematic of the helminth-rabbit system.

Our population has rabbits with no infection (white), single-infections (blue) with either *T. retortaeformis* in the small intestine (black smaller worm) or *G. strigosum* in the stomach (grey larger worm) or dual-infections (red) with both helminths. All rabbits are susceptible to (re-)infection when exposed to infective stages by eating contaminated herbage, starting at around 30 days of age. Species-specific IgA antibodies bind to their targeted helminth species (black and white 'Y'), in our case its excretory/secretory products; in dual infections, specific IgA also reacts with the second helminth through cross-reaction (black and white 'Y' attached to the opposite helminth). Single- and dual-infected animals contribute to the density of free-living stages (black arrows to larvae and eggs on herbage). Rabbits are at risk of infection by exposure through contaminated herbage (white arrows to rabbits). The size and survival of free-living stages is affected by within-host parasite processes and host demography affect, including the direct impact of temperature and humidity. Parameters definition is summarized in table 1.

#### 1.2 Statistical analysis

Statistical analyses were performed using the software R (v.4.0.0) on the following sets of data:

i- model predictions from simulations on population A, i.e. intensity of infection (IOI), shedding (Shed) and risk of infection (RI);

ii- empirical data from population A, i.e. intensity of infection;

iii- empirical data from population B, i.e. intensity of infection and specific IgA.

Generalized Linear Models (GLM) were used to examine changes in the response variable (e.g. IOI) between single and dual infections (SI-DU), entered as categorical variable, and by host age included as continuous explanatory variables. We tested the additive effect of the two variables (SI-DU+Age) as well as the two-way interactions between them (SI-DU∗Age), using individual data and not population averages. We considered GLMs with negative binomial or normal error distributions, based on the type of response variable. This approach was independently performed using parasite and host variables as reported at points *i* to *iii*. We note that in the empirical datasets from population A and B we do not have rabbits of age 1 (i.e. up to 1 month old), which are still un-weaned and not yet infected. Statistical analyses only used available data that start from rabbits of age 2. This is in contrast with figures that report the data by host age, which include age 1 and force the plots to start at age 0. Statistical results are summarized in tables and presented in the supplement.

#### 1.3 Empirical data from population B

Serum IgA against specific excretory/secretory products of adult *G. strigosum* or *T. retortaeformis* was available for every rabbit from population B (Cattadori et al. 2014). An initial data check removed ambiguous cases such as, old rabbits with no helminths but relatively high species-specific IgA responses. Only a handful of these cases were identified.

A preliminary investigation was carried out to examine patterns of IgA, and related intensities of infection, by host age and between single and dual-infected rabbits. Generalized linear models were applied to IgA Optical Density (O.D.) index data, or intensity of infection (IOI) data, as a response variable and following the approach described in SI-1.2. Specific IgA significantly increased with rabbit age and reached an asymptotic trend for *T. retortaeformis*, while steadily increased for *G. strigosum* (figure S2*a* and *b*, table S1). For both helminths, IgA levels were significantly higher in dual- than single-infected rabbits, the two-way interaction SI-DU∗age was not significant and was excluded from the model. Intensities of infection well matched these patterns for both helminths, specifically, intensities were significantly higher in dual- than single-infected rabbits, including the two-way interaction with age (figure S2*c* and *d*, table S1).

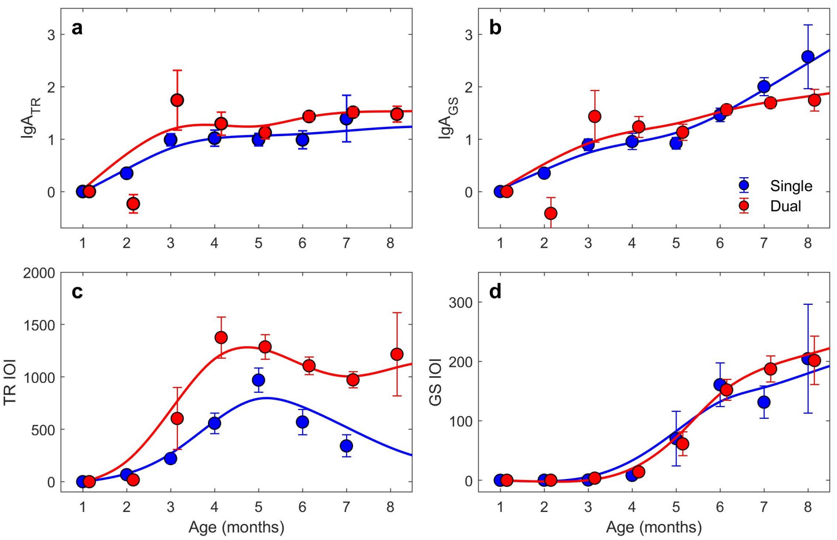


**Figure S2**. Relationships between IgA Optical Density (O.D.) index response (a and b) or intensity of infection (IOI) (c and d) by host age for *T. retortaeformis* (TR, a and c) and *G. strigosum* (GS, b and d), in single- (blue) and dual-infected (red) host groups in population B. A 4$^{th}$ order polynomial curve, weighted by sample size, is fitted to the mean IgA levels or the mean intensities of infection, where mean infection intensities are calculated under the assumption that they follow a negative binomial distribution. The intensity of infection at age 1 month is forced to start at 0 based on the assumption that rabbits are exposed to the risk of infection from about 30 days of age (age class 2). The mean and S.E. are reported. Note that the O.D. index can generate negative values when the IgA response is extremely low (Murphy et al. 2011). Small S.E. are masked by the circles.

These age-related trends are consistent with the general scenario of rabbits constantly exposed to free-living infective stages that lack the ability to develop a robust, long-term protection to reinfections, despite the constant stimulation of an IgA response and, reasonably, many other immune components (Cattadori et al. 2019).

The dynamics of the two helminths from population B exhibited trends similar to the empirical data on infection from population A (compare figure S2 with figure 2 main text and related table S1 with table S3 and S4). This contributes to support the use of IgA data from population B to inform the parameterization of IgA for population A (more in the main text).

**Table S1.** Generalized linear models (with errors following a normal distribution for IgA or a negative binomial distribution for IOI) for the relationships in figure S2 based on individual data. Rabbit age is included as a continuous variable and SI-DU as a factor. The model is fitted to available data that have no age 1 rabbits. The coefficients, Standard Errors, number of observations and related p-values are reported. SI-DU= single-dual infection, Age∗SI-DU= two-way interaction.

| **Response** | **Independents** | **Coefficient [±SE]** | **p-value** |
|---|---|---|---|
| *TR IgA* | Age | 0.089±0.033 | 0.0087 |

| | | | |
|---|---|---|---|
| *(Fig. S2a)* | SI-DU | 0.296±0.117 | 0.0115 |
| | No. obs. | | 1200 |
| *GS IgA (Fig. S2b)* | Age | 0.167±0.039 | <0.00001 |
| | SI-DU | 0.379±0.155 | 0.0143 |
| | No. obs. | | 977 |
| *TR IOI (Fig. S2c)* | Age | 0.544±0.048 | <0.00001 |
| | SI-DU | 3.495±0.292 | <0.00001 |
| | Age∗SI-DU | -0.613±0.061 | <0.00001 |
| | No. obs. | | 1200 |
| *GS IOI (Fig. S2d)* | Age | 1.604±0.075 | <0.00001 |
| | SI-DU | 13.432±0.485 | <0.00001 |
| | Age∗SI-DU | -1.774±0.085 | <0.00001 |
| | No. obs. | | 977 |

#### 1.4 Structure of host populations A and B

The dynamics of the host populations A and B exhibited a clear seasonal pattern that was primarily driven by the annual recruitment of newborns, mostly between April and July (figure S3). The juvenile profile followed that of kittens (age 2 and 3 months) but with a lag of a couple of months, while the number of adults remained relatively high throughout the year in population A, and less so in population B. Population B covered a shorter period of time and was represented by a smaller sample size than population A, and thus more variability in the trends, however, the three main age groups exhibited similar general patterns indicating that the two nearby populations have similar demography. Additional details on the structure and breeding phenology of population A are reported in Mignatti et al. (2016).

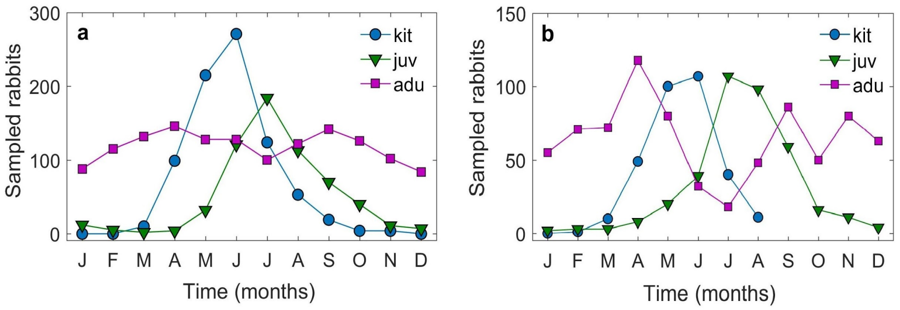


**Figure S3.** Changes in the total number of kittens, juveniles and adults collected in population A (annual range: 1980-2002, a) and population B (2005-2010, b) every month.

A further investigation of population A showed that the proportion of dual-infected rabbits sampled every month was consistently high and between 0.6 and 0.9 for both helminths. There was an exception for *T. retortaeformis* during the May to August period when most of the young and newborn rabbits were circulating and carried single infections (figure S4).

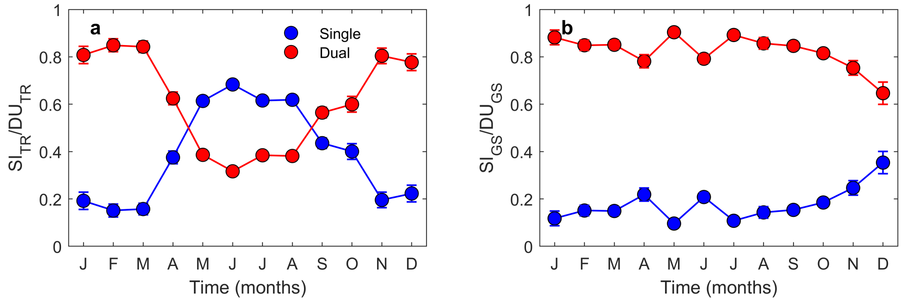


**Figure S4.** Changes in the relative number of rabbits sampled in population A with single and dual infections of *T. retortaeformis* (TR, a) and *G. strigosum* (GS, b) every month.

#### SI-2 Methods

##### 2.1 Newborn recruitment in population A

Our immune-epidemiological model includes the demography of the rabbit population, which is represented by birth and death processes. We modelled the seasonal distribution of newborn recruitment, *f*(*t*), in population A as a beta probability density function calibrated against the available sampled fraction of 2-months old kittens, the age when rabbits start feeding on herbage and get infected with parasites (figure S5). As already noted, and well captured by the beta probability function, most of the newborns circulate in between March and June. The natural mortality rate of the host ($\mu_H$ = 0.0069 days$^{-1}$) was based on a survival of 8.2% for females up to the age of one year under normal conditions (Smith and Trout 1994).

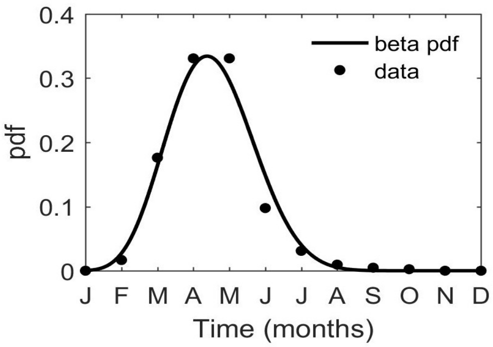


**Figure S5.** Host recruitment function *f*(*t*) modelled as probability density function (pdf) of a beta distribution (line) fitted to the fraction of 2-months old rabbits (circles) by sampling month from population A.

#### SI-3 Results

##### 3.1 Model selection on population A

Among the competing models described in table 2, model selection indicated that for both helminths M2 (specific immunity+intensity-dependence) and M4 (specific immunity+cross-immunity+intensity-dependence) were selected for single and dual infection, respectively (table S2). This indicates that host immunity and parasite demographic processes are both important for the regulation of the intensity of infection of the two helminths. In dual infected rabbits, the unequal cross-immunity also contributes to explain the mechanism of interaction between the two species.

**Table S2.** Model selection summary between competing models M0-M4. $\Delta AIC$ is reported for *T. retortaeformis* (*TR*), *G. strigosum* (*GS*), in single (SI) and dual (DU) infections.

| Models | $\Delta AIC$ ***TR*-SI** | $\Delta AIC$ ***TR*-DU** | $\Delta AIC$ ***GS*-SI** | $\Delta AIC$ ***GS*-DU** |
|---|---|---|---|---|
| M0 | 1,103,660 | 23,043 | 299 | 837 |
| M1 | 1,075,260 | 14,632 | 72 | 804 |
| M2 | 0 | 16 | 0 | 52 |
| M3 | - | 12,513 | - | 741 |
| M4 | - | 0 | - | 0 |

##### 3.2 Annual intensities of infection of population A

The estimated time series of multi-annual infection, represented as 3-month moving average, did describe the general intra-annual variation in parasite infection but missed to capture some of the larger variation observed, particularly for *T. retortaeformis* dual infection (figure S6).

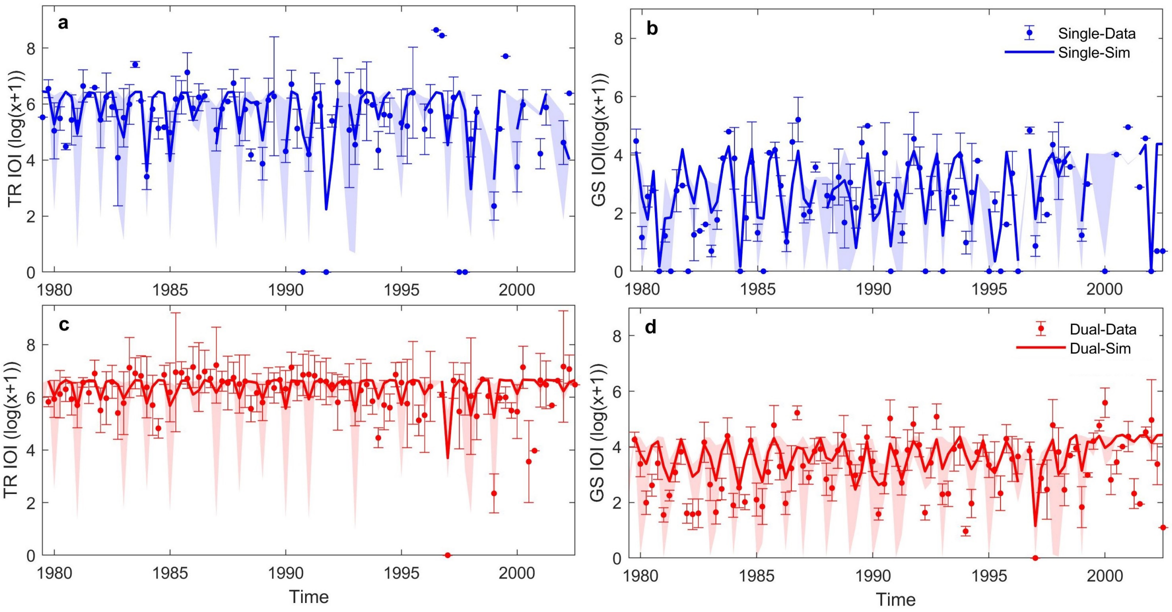


**Figure S6.** Intensity of infection (IOI) log-transformed for *T. retortaeformis* (TR, a and c) and *G. strigosum* (GS, b and d) in single- (a and b, blue) and dual-infected (c and d, red) host groups from population A. The predicted (lines) and empirical (points) monthly time series are reported. The model was fitted to individual data and 3-month averages are presented with the model confident intervals (shaded bands) or the S.E. (bars) for the empirical data.

##### 3.3 Intensities of infection and IgA response of population A

Here we report on the statistical results that support model simulations and empirical data presented in figure 2 (table S3) and figure 4 (table S4). The general description of the statistical analysis is reported in SI-1.2 while the detailed presentation of the results is included in the main text. Both simulated and empirical intensities of *T. retortaeformis* were significantly higher in dual than single infections, the two-way interaction with age was also significant (table S3). Similarly, *G. strigosum* intensities of infection, from simulated and empirical data, were significantly higher in dual than single infections, including the two-way interaction with age (table S3).

**Table S3** Generalized linear models (GLM) for the relationships of *T. retortaeformis* and *G. strigosum* in figure 2, based on individual data from simulations (simul.) and empirical records (empir.). Rabbit age is included as a continuous variable and SI-DU as a factor. The model is fitted to available data that have no age 1 rabbits. The coefficients, Standard Errors, number of observations and related p-values are reported. SI-DU= single-dual infection, Age∗SI-DU= two-way interaction.

| Response | Independents | Coefficient±SE | p-value |
|---|---|---|---|
| *TR IOI simul. (Fig. 2a)* | Age<br>SI-DU<br>Age∗SI-DU | 0.549±0.016<br>1.970±0.016 | <0.00001<br><0.00001<br><0.00001 |

| | | | |
|---|---|---|---|
| | No. obs | -0.316±0.025 | 1977 |
| *TR IOI empir.* *(Fig. 2a)* | Age | 0.387±0.024 | <0.00001 |
| | SI-DU | 1.910±0.205 | <0.00001 |
| | Age∗SI-DU | -0.295±0.037 | <0.00001 |
| | No. obs | | 1977 |
| *GS IOI simul.* *(Fig. 2b)* | Age | 0.359±0.058 | <0.00001 |
| | SI-DU | -1.762±0.401 | <0.00001 |
| | Age∗SI-DU | 0.337±0.060 | <0.00001 |
| | No. obs | | 1325 |
| *GS IOI empir.* *(Fig. 2b)* | Age | 0.232±0.106 | 0.0288 |
| | SI-DU | -1.977±0.734 | 0.0070 |
| | Age∗SI-DU | 0.294±0.111 | 0.0079 |
| | No. obs | | 1325 |

The simulated IgA responses ($r_{ii}I$) presented in figure 4 of the main text were also examined using GLMs as described in SI-1.2. Specific IgA against *T. retortaeformis* was significantly lower in dual than single infections and significantly increased with host age (table S4). In contrast, the specific IgA response against *G. strigosum* was significantly higher in dual than single infections, and also significantly increases with age (table S4). It is important to note that despite these significant patterns the response to *G. strigosum* remains consistently low (figure 4*b*).

**Table S4.** GLMs for the relationships of specific IgA ($r_{ii}I$) to *T. retortaeformis* and *G. strigosum* in figure 4 of the main text, based on individual data from simulations. The model is fitted to available data that have no age 1 rabbits. The coefficients, Standard Errors, number of observations and related p-values are reported. SI-DU= single-dual infection.

| Response | Independents | Coefficient±SE | p-value |
|---|---|---|---|
| $r_{TT}$IgA *(Fig. 4a)* | Age | 0.046±0.066 | 0.0032 |
| | SI-DU | -0.292±0.054 | <0.00001 |
| | No. obs | 1200 | |
| $r_{GG}$IgA *(Fig. 4b)* | Age | 0.029±0.008 | 0.0003 |
| | SI-DU | 0.162±0.031 | <0.00001 |
| | No. obs | 977 | |

##### 3.4 Parasite shedding and Risk of infection by population A

The estimated rate of shedding, $\alpha'_{pi}$, assumes that the amount of eggs shed in the environment is proportional to the intensity of infection, and considers: the total number of eggs shed by an adult parasite per unit of time, $s_i$, the survival of free-living stages shed by a single host, $\alpha_{pi}$, and the rate of total annual recruitment of the host population, $R$. From here, the degree of shedding is calculated as: $Shed_i(a,t) = \alpha'_{pi}\, e^{-\mu_H a} f(t-a) P_i(a,t)$. Hence, the load of eggs shed is the outcome of the interaction between parasite reproduction and the age-dependent dynamics of the host population, where $e^{-\mu_H a}$ is host survival and *f*(*t*) the temporal distribution of host reproduction. To reduce model complexity, we assume that eggs directly hatch into infective larvae and the whole population of free-living stages (eggs and larvae) is directly affected by temperature and humidity.

Simulations showed that for both helminths, the mean number of eggs shed was significantly higher in dual- than single-infected rabbits when host age were considered, a trend particularly clear for *T. retortaeformis* (figure S7, table S5). The negative two-way interaction SI-DU∗age is probably caused by the decrease of shedding with host age for *T. retortaeformis*, and the increase of shedding in older single-infected host for *G. strigosum.*

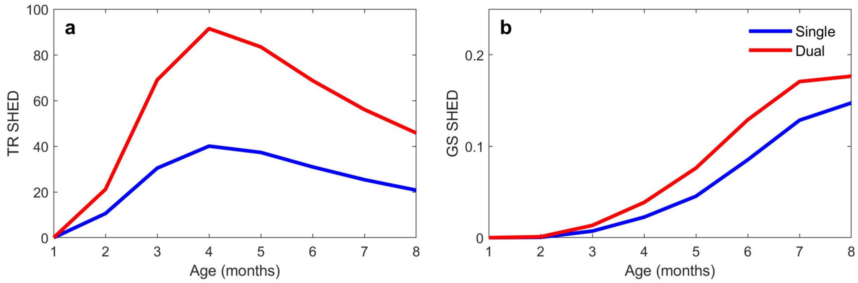


**Figure S7.** Estimated mean shedding (SHED) log-transformed by host age for *T. retortaeformis* (TR, a) and *G. strigosum* (GS, b) in single- (blue) and dual-infected (red) host groups from population A.

**Table S5.** GLMs for the relationships in figure S7 for *T. retortaeformis* (TR) and *G. strigosum* (GS). The model is fitted to available data that have no age 1 rabbits. The coefficients, Standard Errors, number of observations and related p-values are reported. SI-DU= single-dual infection, Age∗SI-DU= two-way interaction.

| Response | Independents | Coefficient [±SE] | p-value |
|---|---|---|---|
| *TR Shed (Fig. S7a)* | Age | -0.033±0.004 | <0.00001 |
| | SI-DU | 0.807±0.027 | <0.00001 |
| | Age∗SI-DU | -0.002±0.005 | <0.00001 |
| | No. obs. | | 134304 |
| *GS Shed (Fig. S7b)* | Age | 0.385±0.008 | <0.00001 |
| | SI-DU | 0.648±0.065 | <0.00001 |
| | Age∗SI-DU | -0.059±0.010 | <0.00001 |
| | No. obs. | | 134304 |

Our modelling approach allows us to estimate what group of hosts (single- or dual-infected) contributes the most to the production of these free-living stages and, in turn, transmission.

Simulations indicated that throughout the months, rabbits dual-infected generated significantly more viable free-living stages of both helminths, than single-infected rabbits (figure 5, table S6).

**Table S6.** GLMs for the relationships of figure 5 for *T. retortaeformis* (TR) and *G. strigosum* (GS). The coefficients, Standard Errors, number of observations and related p-values are reported. SI-DU= single-dual infection, Month∗SI-DU= two-way interaction.

| Response | Independents | Coefficient [±SE] | p-value |
|---|---|---|---|
| *TR RI* | Month | 0.165±0.006 | <0.00001 |
| | SI-DU | 1.724±0.059 | <0.00001 |
| *(Fig. 5a)* | Month∗SI-DU | -0.015±0.008 | <0.00001 |
| | No. obs. | | 16790 |
| *GS RI* | Month | 0.130±0.005 | <0.00001 |
| | SI-DU | 1.734±0.054 | <0.00001 |
| *(Fig. 5b)* | Month∗SI-DU | -0.078±0.007 | <0.00001 |
| | No. obs. | | 16790 |

**References**


Cattadori et al. 2014. *Ecology* 95, 1684-1692.
Cattadori et al. 2019. *Ecol. Evol.* 9, 13495-13505.
Mignatti et al. 2016. *PNAS* 113, 2970-2975.
Murphy et al. 2011. *Paras. Immunol.* 33, 287-302.
Smith and Trout 1994. *J. Appl. Ecol*. 31: 223-230.
